# Suitability of clove oil for the euthanasia of four ornamental fish species: Goldfish (*Carassius auratus*), Bristlenose catfish (*Ancistrus* sp.), Midas cichlid (*Amphilophus citrinellus*) and Three-spot gourami (*Trichopodus trichopterus*)

**DOI:** 10.64898/2026.10.31.763359

**Authors:** Livia Brülhart, Lara Czichelski, Gary Delalay, Nicolas Diserens

## Abstract

Euthanasia of ornamental fish is usually performed by immersion in an anaesthetic bath. However, it may be difficult for private fish owners to access anaesthetics, and alternative methods are limited. Clove oil is commonly available in drugstores and its use for fish euthanasia has been reported. However, there is a lack of scientific literature supporting its suitability for this purpose and of available species-specific protocols. Our study aimed to assess the suitability of clove oil to euthanise goldfish (*Carassius auratus*), bristlenose catfish (*Ancistrus* sp.), Midas cichlid (*Amphilophus citrinellus*) and three-spot gourami (*Trichopodus trichopterus*). Time to loss of reflexes and to cessation of opercular movements were recorded. Aversive reactions and histological gill lesions were compared with positive and negative controls, and reversibility of euthanasia attempts was assessed. Opercular movements did not reliably cease in goldfish, even under high clove oil concentrations (1000 mg/L). Clove oil caused loss of indicators in Midas cichlid at a concentration of 1000 mg/L, and in bristlenose catfish and three-spot gourami at a concentration of 400 mg/L. Five Midas cichlid recovered after treatment with clove oil. Clove oil caused severe gill lesions to goldfish and Midas cichlids. The results show that clove oil is not suitable for humane euthanasia of goldfish and Midas cichlid. Additionally, some studies have questioned the ability of clove oil to induce loss of consciousness in fish. Therefore, we recommend avoiding euthanasia with clove oil when suitable alternatives are available and using a secondary killing method, as euthanasia attempts are not always irreversible.

## Introduction

Euthanasia is defined by Yeates (2010) as the killing of an animal in the animal’s interest by a human who believes it is in the animal’s interest. This definition implies that euthanasia is an important tool for animal welfare but is limited to situations in which the animal itself benefits from being killed, such as when it is suffering from disease and therapy is not an option. Ornamental fish are also sometimes sacrificed for other purposes which, although not directly in the animal’s interest, may still be ethical. For instance, sacrificing fish for population management can be ethical, as it is often not possible to limit reproduction through other ways and it allows to preserve the welfare of the remaining fish (DiVincenti et al. 2023; Leary et al. 2020). Additionally, sacrificing some fish to perform a pathological examination can allow for rapid and effective diagnosis of the cause of a disease, thereby enabling treatment of the remaining population (Ross 2001). Regardless of whether such killing can be classified as euthanasia, these practices are considered ethical and important for ensuring welfare at the population level. For the sake of simplicity, the term euthanasia will be employed to refer to the ethical sacrificing of fish throughout the manuscript, without further differentiation based on its purpose.

The term euthanasia derives from the Greek words for “good death”, implying that the process should be humane, minimizing or eliminating pain and distress (Leary et al. 2020). This highlights the importance of reproducible and consistent methods. In ornamental fish, euthanasia is most often performed through immersion in an anaesthetic bath (Neiffer and Stamper 2009; Leary et al. 2020), and the drug should, thus, minimize aversive responses and adverse effects in fish while ensuring that loss of consciousness is induced as quickly as possible.

Classical anaesthetics used for euthanasia of ornamental fish comprise, among others, buffered tricaine, benzocaine and phenoxyethanol (Neiffer and Stamper 2009; Smith and Harms 2023; Leary et al. 2020). However, these are classified as drugs in many countries and therefore not always easily available to private owners of ornamental fish. In Switzerland for example, no drugs are registered for the anaesthesia and euthanasia of fish. Therefore, they are available exclusively to veterinarians, who are the only ones allowed to import it from abroad (article 7 of the *Tierarzneimittelverordnung* [Swiss Ordinance on Veterinary Medicinal Products], Swiss Federal Council 2025). However, in practice, only a handful of specialized fish veterinarians do import such drugs, which renders them relatively inaccessible to private fish owners. Consequently, some owners may refrain from euthanising their fish, potentially harming their welfare, or may release fish from overpopulated ponds or aquaria into the wild, with potential serious welfare and ecological implications (Banha et al. 2019; Padilla and Williams 2004; Gertzen et al. 2008).

Given that physical methods, such as stunning caused by blunt force followed by death through exsanguination, are often not considered acceptable by fish owners (Leary et al. 2020) or are impractical due to the size of the fish, there is a need for a suitable alternative. To this end, establishing protocols for the use of substances not categorized as drugs and which would be easily accessible for both private owners and non-specialized veterinarians is of utmost importance.

Clove oil, whose main compound is eugenol, is not classified as a drug in Switzerland and can easily be purchased from drugstores. Nevertheless, eugenol does have pharmacological effects. It is commonly employed as an anaesthetic in fish (Smith and Harms 2023; Endo et al. 1972; Wallbom et al. 2026; Stetter 2001; Neiffer and Stamper 2009; Hikasa et al. 1986), and its use has been described as a humane method to euthanise fish (Smith and Harms 2023; Leary et al. 2020; Von Krogh et al. 2021; Fernandes et al. 2016). However, its efficiency on all commonly kept ornamental fish is still to be demonstrated, and ideal species-specific dosages are still to be evaluated and reported. Indeed, there is limited scientific literature supporting the suitability of eugenol for the humane euthanasia of ornamental fish, as well as a lack of available species-specific protocols.

Therefore, this study aimed to assess the suitability of eugenol to euthanise different freshwater species of aquarium and pond fish and, if found suitable, to determine the ideal dosage that could be used by private owners to euthanise their fish.

## Material & Methods

### Overview

We performed three sequential experiments on four different fish species to determine if the use of clove oil is suitable to euthanise fish from these species.

The first experiment consisted of comparing the effects of up to four different eugenol concentrations per species, allowing for the determination of ineffective and harmful concentrations. Based on these results, we could identify the potential optimal eugenol concentration for euthanasia of each species.

The second experiment, based on the results obtained through the first experiment, aimed to assess the reliability of the optimal concentration identified for each species. Following exposure to treatment and both loss of reflexes and disappearance of opercular movements, fish were transferred to a recovery tank for an hour to evaluate whether they could recover.

The third experiment focused on evaluating the potential adverse reactions to eugenol. We compared changes in behaviour and presence of gill lesions in fish treated with eugenol to those in untreated fish kept in plain water (negative control) and in fish treated with buffered tricaine methanesulfonate (positive control). Buffered tricaine (Tricaine Pharmaq 1000 mg/g, PHAR-MAQ AS, Overhalla, Norway) was chosen as a positive control, as it is the drug most widely used to euthanise fish in the European Union.

### Origin of the animals and detention

The study focused on four species: goldfish (*Carassius auratus*), bristlenose catfish (*Ancistrus* sp.), Midas cichlids (*Amphilophus citrinellus*) and three-spot gouramies (*Tri chopodus trichopterus*). These fish species were chosen for the following reasons: 1) they are some of the most common species of pet fish in Switzerland; 2) they are kept in both aquarium and ponds; 3) they are phylogenetically distant from each other, and 4) they cover a wide range of fish sizes. Moreover, the inclusion of three-spot gourami in this study allowed us to test the euthanasia method on an anabantoid species, which are characterised by possessing a lung-like labyrinth organ which allows them to capture oxygen from the air, temporarily pausing branchial breathing (Tate et al. 2017). Furthermore, the inclusion of goldfish allowed us to test a clove oil protocol on a fish species which is known to be difficult to euthanise using chemical agents (Balko et al. 2018; Perret-Thiry et al. 2022; Heniff and Balko 2023). Further criteria for species selection were: the avoidance of using wild caught fish; and the preference for including fish that would nevertheless have been euthanised as part of population management due to overpopulation issues. The goldfish in this study originated from two private holdings dealing with overpopulation issues. The bristlenose catfish and the Midas cichlids were provided by the zoological garden Papiliorama Foundation (Kerzers, Switzerland) and were due to be euthanised as part of population management. Because of the lack of alternatives, the three-spot gourami were obtained from four different fish and pet shops. Healthy adult fish, exhibiting no signs of external injury or disease, were selected for this study and randomly assigned to a group in one of the three experiments of the study.

The bristlenose catfish and the three-spot gourami were housed indoor in a flow-through system composed of 130 L freshwater tanks, with maximum densities of 40 and 10 individuals per tank, respectively. The tanks were continuously aerated with air stones, contained artificial plants and were obscured from one side. Additionally, hiding places in the shape of small plant pots made of clay were added. The fish were held in a 14:10 day-night regime, with light from 06:00 to 20:00 and darkness from 20:00 to 06:00. Water quality and fish health were assessed daily. Daily water quality assessments were conducted following standardised protocols, including temperature (24.5 °C – 27 °C), total ammonia nitrogen (≤ 0.2 mg/L) and oxygen (≥ 7.5 mg/L), using standard test kits (Merck – Germany and OxyGuard – Denmark, respectively). Nitrite and pH were only measured when deemed necessary. The three-spot gourami were fed commercial fish feed (TetraMin Flakes, Tetra GmbH, Melle, Deutschland) every day. The bristlenose catfish received special catfish food (Sera Wels-Chips, Sera GmbH, Heinsberg, Deutschland) and a slice of cucumber daily.

The goldfish were maintained in groups of maximum 21 fish in a semi-outdoor environment under a 12:12 dark/ light regime. The 2000 L fish tanks were supplied with an aeration pump and were enriched with shelters consisting of branches and stones. Water quality and fish health was assessed daily, and water changes were performed as needed to maintain the optimal water quality parameters. Water quality assessments were performed as described for bristlenose catfish and three-spot gourami. Tanks were filled with municipal tap water with a pH of 7.6, total ammonia nitrogen concentration remaining at 0.5 mg/L or below (excepting an isolated peak at 1.5 mg/L), and the oxygen saturation ranging from 74% to 93%. Water temperature varied with weather conditions and ranged from 10.1 °C to 24.3 °C. Fish were fed a commercial pelleted diet (Tetra Pond Sticks, Tetra GmbH, Melle, Germany).

The Midas cichlid were kept by the Papiliorama Foundation under their usual husbandry conditions until the beginning of the experiment.

All fish were acclimated for a minimum of two weeks prior to experimentation, with the exception of the Midas cichlid, as the trials were performed directly at their housing site, and were determined to be clinically healthy based on repeated visual examinations.

The distribution of fish size and weight per species for each treatment, concentration and experimental step can be found in Appendix 1.

### Study design

#### First experiment: determination of the effective con centration

The aim of the first experiment was to determine the lowest effective dose of eugenol, and the hypothesis was that this dose would also cause the fewest lesions and the least adverse reactions. Three-spot gouramies were exposed to eugenol concentrations of 170, 250 and 400 mg/L, bristlenose catfish to concentrations of 250 and 400 mg/L, and goldfish and Midas cichlids to concentrations of 250, 400, 650 and 1000 mg/L, according to available literature (Leary et al. 2020; Ayala-Soldado et al. 2023; Fernandes et al. 2016; Day et al. 2024).

Each dose was tested with up to three fish per species and was considered effective if loss of righting reflex (LoRR), loss of startle reflex (LoSR) and cessation of opercular movement (CoOM) occurred for all tested fish within the time frame defined in the humane end-points. If absence of righting reflex, startle reflex and opercular movements persisted for 60 seconds, a gill cut or decapitation was performed to ensure death. Histological samples were taken from the gills and evaluated for lesions and pathological changes.

#### Second experiment: assessment of reversibility of eu thanasia attempt

The first experiment provided us a potentially optimal eugenol concentration for euthanasia of bristlenose catfish, three-spot gouramies and Midas cichlids. The reliability and effectiveness were then evaluated in the second experiment by exposing up to thirty fish per species to this potentially optimal eugenol concentration. We did not perform this step for goldfish, as we could not determine a reliable effective dose in the first experiment.

Ten minutes after LoRR, LoSR and CoOM, fish were transferred to an aerated recovery tank filled with water collected from their original tank. There, we monitored the fish over one hour to assess any signs of consciousness, such as the presence of a righting reflex or opercular movements. If the fish did not display signs of consciousness after an hour, startle reflex was assessed, and, if negative, reaction to deep pain was evaluated, by firmly compressing the caudal peduncle with forceps (Stetter 2001).

Fish that showed no response to these reflex tests were considered dead, and a gill cut or decapitation was performed as a secondary euthanasia method, as recommended by Leary et al. (2020). If a fish showed signs of consciousness, the trial was stopped immediately, and the fish was euthanised using buffered tricaine followed by gill cut or decapitation.

If the concentration previously defined in the first step failed to provoke LoRR, LoSR or CoOM in a fish within the accepted time frame, the second experiment was interrupted and restarted using a higher dosage.

#### Third experiment: assessment of adverse effects

In the third experiment, we compared the effects of three treatments on the different species:

a. clove oil at the optimal concentration determined during experiments 1 and 2;
b. a positive control with buffered tricaine at 600 mg/ L for bristlenose catfish and three-spot gourami, and 1000 mg/L for Midas cichlids; and
c. a negative control with water from their original aquarium. As for the second experiment, goldfish were not included in this step, as the first experiment did not allow us to determine an effective dose for this species.

Twenty fish per species were exposed to each treatment group. Treatments a) and b) were compared to evaluate the differences in pathological effects between clove oil and tricaine. The negative control group was included to differentiate stress behaviour caused by the treatment and by handling alone. As soon as LoRR, LoSR and CoOM were sustained for 60 seconds in fish from the groups a) and b), a gill cut or decapitation was performed to ensure death and histological samples were taken.

To reduce the number of animals, in accordance with the 3Rs principles, we first performed exposure to negative control (i.e. water only) and reused the same fish again for exposure to treatment (eugenol) or to positive control (buffered tricaine). A minimum of a 24-hour break was respected between exposure to negative control and to treatment and positive control to allow fish to recover from handling stress.

#### Ethical considerations and humane endpoints

The experiments were interrupted if a fish did not lose righting reflex within 10 minutes or if opercular movements did not cease within 20 minutes. If an experiment was interrupted, the dosage was deemed inadequate and the experiment was reconducted with a higher dosage.

The project followed the regulations of the Swiss Animal Welfare Act (SR 455) and Swiss Animal Welfare Ordinance (SR 455.1). All experiments were approved by the Cantonal Veterinary Office Bern, Switzerland under Authorization #BE50/2024.

### Treatment

Clove oil (84.4% Eugenol; Hänseler Swiss Pharma®, Herisau, Switzerland; see Appendix 2 for the certificate of analysis) was mixed with ethanol (100%) in a 1:9 ratio to ensure water solubility, as clove oil alone is poorly soluble in water (Smith and Harms 2023). The required solution was prepared as a stock solution. In the first experimental step, the clove oil solution was changed after each animal. In the second and third experimental steps, the solution was changed after every fifth animal for efficiency and economic reasons. As the eugenol content of clove oil was known, concentrations were calculated for eugenol directly, and not for clove oil.

The tricaine solution used in the positive control group was buffered with 5 mL of sodium bicarbonate (NaHCO_3_) per litre of tank water to maintain an appropriate pH.

Exposure was conducted in 2 L tanks for bristlenose catfish and three-spot gourami, 4 L tanks for goldfish, and 10 to 15 L tanks for the Midas cichlids, depending on fish size. The water was taken directly from the tanks the fish were housed in. Water temperatures ranged from 14.2 to 19.4 °C for goldfish, 19.9 to 22.8 °C for Midas cichlid, 20.3 to 24.5 °C for bristlenose catfish, and 21.9 to 25.0 °C for three-spot gourami (Appendix 3). Clove oil and tricaine, respectively, were dissolved at the target concentration in the experimental tank before the fish were transferred into it. Each fish was selected randomly and exposed to treatment individually.

### Measures of indicators

Following the immersion of the fish in the treatment tank, righting reflex and opercular movements were monitored. LoRR and CoOM were visually determined, without intervention from the observer. For bristlenose catfish, CoOM was replaced with arrest of breathing (which, for sake of simplicity, will also be reported as CoOM throughout the manuscript) due to their different anatomy (Geerinckx and Adriaens 2006). To this end, the fish were turned on their backs after LoRR, so that breathing could be monitored by observing buccal movements associated with respiration.

After LoRR, startle reflex was triggered every 15 seconds by dropping a 76 g metal marble on the table from a height of 15 cm next to the tank until the fifth minute, and then every minute thereafter. The time taken until LoRR, LoSR and CoOM were separately documented and video material was recorded.

The experiment was terminated if at least one of LoRR, LoSR, or CoOM did not occur within twenty minutes, or if any of the following criteria were met: tachypnoea or frantic swimming persisting for more than 30 seconds; or jumping or tremors lasting longer than 15 seconds. In such cases, the fish were euthanised using buffered tricaine, followed by a gill cut or decapitation.

### Behavioural observations

During induction, each fish was individually observed by an experimenter, and video material was recorded. Particular attention was given to stress indicators such as fast swimming, atypical swimming patterns, gasping air at the surface, and atypical behaviour. Based on the observations and the video material, differences between treatments were evaluated qualitatively and subjectively. No blinding was applied.

### Histology

In the first and in the third experiments, whole euthanised bristlenose catfish, three-spot gourami and goldfish as well as gills sampled from the euthanised Midas cichlids were fixed in 10% buffered formaldehyde for 24 hours. Fixed tissues were then decalcified with Kristensen’s solution (136 g Na-Formiate (Merck KGaA, Darmstadt, Germany) and 332 mL furmic acid 98% (Merck KGaA, Darmstadt, Germany) diluted in 4 L distilled water) for 12 to 24 hours. The gills were then embedded in paraffin, and sections were cut at 4 µm and stained with hematoxylin-eosin (H&E). For each histological cut, three random fields were examined under a light microscope at 200x magnification for a) epithelial lifting; b) oedema; c) epithelial proliferation; d) aneurisms in the lamellaes; and e) presence of parasites, and at 400x magnification for f) presence of eosinophilic granular cells. Epithelial lifting, oedema and aneurysms were hypothesized to be relevant and potentially treatment-related, whereas epithelial proliferation, the presence of parasites, and the presence of granular cells were assessed as quality-control parameters to exclude other potential causes of differences in gill lesions between groups.

On average, three gill arches per fish were analysed. A semi-quantitative evaluation was performed, and the histopathological changes were evaluated and classified according to Table 1. Each field was scored separately, and data was then aggregated at fish level by calculating the mean of all three fields for each assessed parameter. The gills of one Midas cichlid in the clove oil group could not be processed for technical reasons.

**Table 1.**
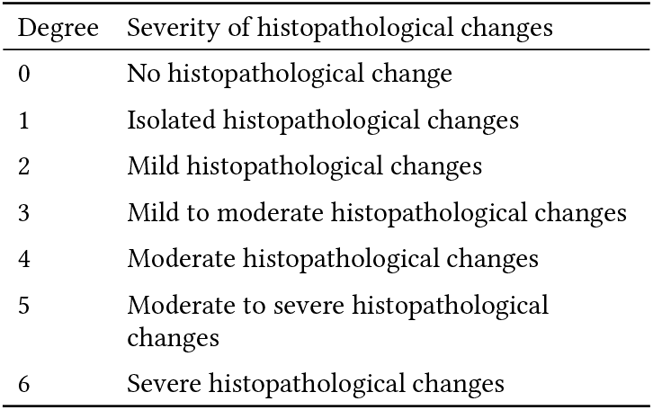
Semi-quantitative histopathological evaluation.

All histological cuts from fish from the same species were assessed by the same scorer to allow comparison between individuals and treatments within a species. However, not all species were assessed by the same scorer, and intraspecific comparisons could, therefore, not be performed. No blinding was applied.

### Statistical analysis

Data was collected in a Microsoft Excel 2023 spreadsheet and imported in R (version 4.6.1, released 2026-06-24) (R Core Team 2025) using the package readxl 1.5.0.1 (Wickham and Bryan 2025). For data manipulation and transformation, the packages dplyr 1.2.1 (Wickham, François, et al. 2026) and tidyr 1.3.2 (Wickham et al. 2025) were used. Visualizations were created with the package ggplot2 4.0.3.9000 (Wickham, Chang, et al. 2026; Wickham 2016).

A logistic regression was performed in R to assess association between fish weight (explanatory variable) and recovery (binary response variable). The model was fitted using a binomial error distribution with a logit link function.

Differences in histological lesion score between treatments were assessed separately for each fish species and type of histological lesion using the Wilcoxon ranksum test. To account for multiple comparisons, p-values were adjusted within each species using the Holm–Bonferroni procedure, treating all three tests performed on fish from the same species as one family of hypotheses. Adjusted P values < 0.05 were considered statistically significant.

### Rehoming of surplus animals

As experimentation involved euthanasia, no fish subjected to experimentation survived the project. During experimentation, we could avoid the detention of more Midas cichlid than required for the experiment, given that fish were collected directly from their population at the zoological garden. We could also avoid surplus bristlenose catfish and three-spot gouramies by buying them in small groups. However, this could not be avoided for goldfish, in particular because we did not expect to be unable to perform steps 2 and 3 on this species. We were, however, able to rehome them by giving them to an experienced private owner.

## Results

### Clove oil concentration needed to reliably provoke LoRR, LoSR and CoOM

Table 2 shows the time to LoRR, LoSR and CoOM across species and clove oil concentration as well as the reliability of the treatment to provoke LoRR, LoSR and CoOM. Figure 1 also includes tricaine treatments.

**Table 2.** Time to loss of indicators for goldfish (*Carassius auratus*), bristlenose catfish (*Ancistrus* sp.), Midas cichlid (*Amphilophus citrinellus*) and three-spot gourami (*Trichopodus trichopterus*) treated with different concentrations of eugenol diluted in the water. LoRR: loss of righting reflex / CoOM: cessation of opercular movement / LoSR: loss of startle reflex / N number of fish tested

| Concentration [mg/L] | N | All fish succesfully lost all indicators | Number of animals with LoRR | Median time to LoRR [s] | Standard deviation for time to LoRR [s] | Number of animals with CoOM | Median time to CoOM [s] | Standard deviation for time to CoOM [s] | Number of animals with LoSR | Median time to LoSR [s] | Standard deviation for time to LoSR [s] |
|---|---|---|---|---|---|---|---|---|---|---|---|
| Goldfish |  |  |  |  |  |  |  |  |  |  |  |
| 250 | 2 | No | 2 | 50 | 14 | 1 | 1470 | – | 0 | – | – |
| 400 | 3 | No | 3 | 32 | 12 | 2 | 459 | 14 | 0 | – | – |
| 650 | 3 | No | 3 | 38 | 7 | 2 | 694 | 337 | 1 | 75 | – |
| 1000 | 3 | No | 3 | 61 | 15 | 2 | 266 | 7 | 3 | 75 | 68 |
| Midas cichlid |  |  |  |  |  |  |  |  |  |  |  |
| 250 | 1 | No | 1 | 37 | – | 0 | – | – | 0 | – | – |
| 400 | 2 | No | 2 | 28 | 6 | 1 | 811 | – | 0 | – | – |
| 650 | 2 | No | 2 | 22 | 1 | 1 | 15 | – | 0 | – | – |
| 1000 | 53 | Yes | 53 | 23 | 15 | 53 | 19 | 10 | 11 | 15 | 4 |
| Bristlenose catfish |  |  |  |  |  |  |  |  |  |  |  |
| 250 | 4 | No | 4 | 66 | 22 | 3 | 500 | 134 | 0 | 110 | – |
| 400 | 53 | Yes | 53 | 53 | 21 | 53 | 355 | 151 | – | 140 | 191 |
| Three-spot gourami |  |  |  |  |  |  |  |  |  |  |  |
| 170 | 20 | Yes | 20 | 54 | 71 | 20 | 527 | 145 | – | 64 | 105 |
| 250 | 3 | Yes | 3 | 56 | 14 | 3 | 500 | 40 | – | – | – |
| 400 | 53 | Yes | 53 | 37 | 72 | 53 | 360 | 149 | – | 85 | 144 |

**Figure 1.**
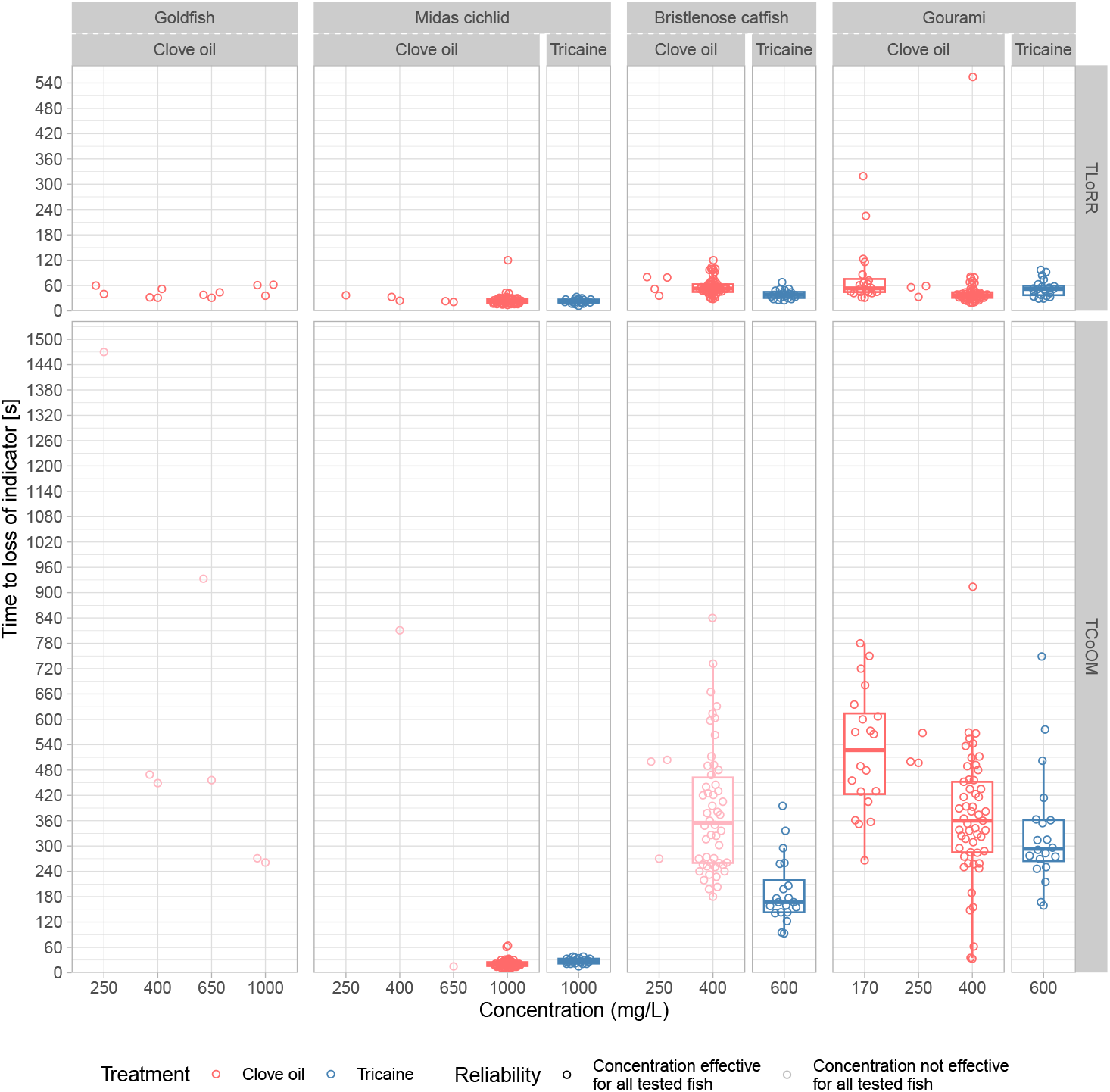
Time to loss of indicators for goldfish (*Carassius auratus*), bristlenose catfish (*Ancistrus* sp.), Midas cichlid (*Amphilophus citrinellus*) and three-spot gourami (*Trichopodus trichopterus*) treated with tricaine and different concentrations of eugenol diluted in the water. TLoRR: time to loss of righting reflex / TCoOM: time to cessation of opercular movement

At all combinations of species and clove oil concentration tested, all fish (n = 204) reliably lost righting reflex within ten minutes.

Opercular movement did not reliably cease in gold-fish, even with high concentrations of eugenol (1000 mg/L).

Startle reflex was not a reliable indicator in our study. In the control group, 5 out of 20 (25%) Midas cichlids did not show startle reflex after the marble test, although they were not anesthetized. Additionally, we incidentally observed that bristlenose catfish could still respond to light finger taps on the table for a considerable time after losing the startle reflex with the marble test. All fish that retained the startle reflex also continued to show opercular movements.

As four three-spot gouramies exposed to 170 mg/L clove oil in experiment 2 exhibited prolonged undulating movements with their fins at the bottom of the aquaria and then surfaced to gasp air, the trial at this concentration was terminated after 17 fish, and the experiment was continued using a concentration of 400 mg/L.

### Reversibility of euthanasia attempts

No bristlenose catfish nor three-spot gourami recovered within an hour after treatment (n = 30 per species). However, 5 out of 30 Midas cichlids recovered. The distribution of body weight for fish who did and fish who did not recover is presented in Figure 2. A logistic regression using fish weight to model reversibility of euthanasia attempt was not significant (p = 0.097).

**Figure 2.**
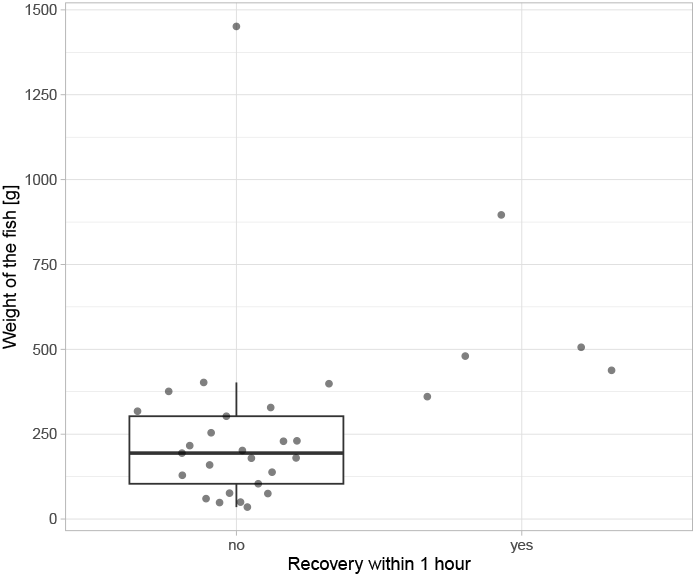
Distribution of Midas cichlid (*Amphilophus cit rinellus*) weight per recovery status. The fish were treated by bath with 1000 mg/mL eugenol and 10 minutes after loss of all reflexes transferred in tank with optimal water condition for maximum one hour. To avoid overplotting, points are jittered horizontally.

### Behavioural observations

Directly after release in the treatment tank, all fish showed signs of stress. Midas cichlid in the negative control group showed the longest and most intense episodes of fast swimming, followed by Midas cichlid euthanised with tricaine. Midas cichlid euthanised with clove oil seldom showed signs of stress in the form of fast swimming but expressed forceful buccal pumping. Midas cichlids euthanised with tricaine also showed forceful buccal pumping similar in form and intensity, but not Midas cichlid in the negative control group. Although signs of stress were observed for bristlenose catfish and three-spot gourami across all treatments after release in the treatment tank, mainly expressed as fast swimming, interaction with the tank wall and release of faeces, no difference in intensity was observed between treatments and no other particular difference in behaviour was observed during induction between treatments. However, five three-spot gourami euthanised with clove oil regained some body and opercular movements after LoRR, loss of movement and CoOM, four of them undulating on the side at the bottom of the tank for a small time before losing both body and opercular movements again. The fifth individual was even able to swim for a short time at the surface of the water, though without regaining the righting reflex. A sixth three-spot gourami treated with clove oil regained movements after LoRR and loss of movements, but before CoOM had occured.

### Histological lesions

Figure 3 shows two examples of severe and mild gill lesions observed in the frame of the experiment. Goldfish showed massive epithelial lifting after clove oil was used, making it impossible to assess oedema and epithelial proliferation for all goldfish but one. This was the case for all dosages used. Midas cichlids also showed high lesion scores, aneurisms and oedema being significantly more present in the gills after treatment with clove oil than after treatment with tricaine, with a median shift of the score of 3.00 and 2.33, respectively. Gourami and bristlenose catfish were less prone to aneurisms and oedema than Midas cichlids but had higher prevalences of epithelial lifting. Contrary to Midas cichlid, three-spot gourami showed significantly less lesions (aneurisms) when treated with clove oil than when treated with tricaine, although the median shift of the score was smaller (-0.67). No significant difference between treatments could be found for bristlenose catfish. Figure 4 shows the distribution of scores for each species, treatment and histological parameter. Table 3 summarises the results of the Wilcoxon rank-sum tests for comparisons between treatment for each species and histological parameter.

**Table 3.** Results of Wilcoxon rand-sum tests for difference in lesion scores in histological examination between clove oil and tricaine euthanasia for goldfish (*Carassius auratus*), midas cichlid (A*mphilophus citrinellus*), bristlenose catfish (*Ancistrus* sp.) and three-spot gourami (*Trichopodus trichopterus*). For all histological parameters, lesions were scored on an integer scale of 0 to 6 at three different locations of the slide, and the mean of the three scores was then attributed to each fish. No tricaine control group was available for goldfish, as the experiment was interrupted as the treatment was not working reliably. To account for multiple comparisons, p-values were adjusted within each species using the Holm– Bonferroni correction, treating all three tests performed on fish from the same species as one family of hypotheses. Provided are the species for which difference was tested, the histological parameter that was considered, the median score and number of animals (N) for each treatment group, the p value both before and after Holm-Bonferroni correction, the W statistic and the Hodges–Lehmann estimate of the location shift, including 95% confidence interval. NA: not applicable

| Parameter | Concentration clove oil / tricaine [mg/L] | Median [N] (clove oil treatment) | Median [N] (tricaine treatment) | p.value | p.adjusted | W | Hodges–Lehmann estimate of the location shift [95% confidence interval] |
|---|---|---|---|---|---|---|---|
| Goldfish |  |  |  |  |  |  |  |
| Epithelial lifting | 250 / NA | 6.00 [1] | NA | NA | NA | NA | NA |
|  | 400 / NA | 5.67 [3] | NA | NA | NA | NA | NA |
|  | 650 / NA | 5.67 [3] | NA | NA | NA | NA | NA |
|  | 1000 / NA | 5.50 [2] | NA | NA | NA | NA | NA |
| Oedema | 400 / NA | 4.67 [1] | NA | NA | NA | NA | NA |
| Aneurism | 250 / NA | 0.33 [1] | NA | NA | NA | NA | NA |
|  | 400 / NA | 0.00 [3] | NA | NA | NA | NA | NA |
|  | 650 / NA | 0.00 [3] | NA | NA | NA | NA | NA |
|  | 1000 / NA | 0.00 [2] | NA | NA | NA | NA | NA |
| Midas cichlid |  |  |  |  |  |  |  |
| Epithelial lifting | 1000 / 1000 | 0.00 [23] | 0.00 [20] | 1.0000 | 1.0000 | 230.0 | 0.00 [ 0.00-0.00] |
| Oedema | 1000 / 1000 | 4.67 [22] | 2.33 [20] | < 0.0001 | < 0.0001 | 425.5 | 2.33 [ 1.67-3.00] |
| Aneurism | 1000 / 1000 | 3.33 [22] | 0.00 [20] | < 0.0001 | < 0.0001 | 407.0 | 3.00 [ 1.67-4.67] |
| Bristlenose catfish |  |  |  |  |  |  |  |
| Epithelial lifting | 400 / 600 | 1.33 [23] | 1.67 [20] | 0.8603 | 1.0000 | 222.5 | 0.00 [−1.00-0.67] |
| Oedema | 400 / 600 | 3.33 [23] | 2.67 [20] | 0.1155 | 0.3464 | 294.5 | 0.33 [ 0.00-1.00] |
| Aneurism | 400 / 600 | 0.00 [23] | 0.00 [20] | 0.6109 | 1.0000 | 247.0 | 0.00 [ 0.00-0.00] |
| Three-spot gourami |  |  |  |  |  |  |  |
| Epithelial lifting | 400 / 600 | 2.67 [23] | 2.83 [20] | 0.9468 | 0.9468 | 233.0 | 0.00 [−0.67-0.67] |
| Oedema | 400 / 600 | 3.00 [23] | 3.00 [20] | 0.4473 | 0.8945 | 198.5 | −0.33 [−1.00-0.33] |
| Aneurism | 400 / 600 | 0.00 [23] | 1.00 [20] | 0.0065 | 0.0194 | 128.0 | −0.67 [−1.00-0.00] |

**Figure 3.**
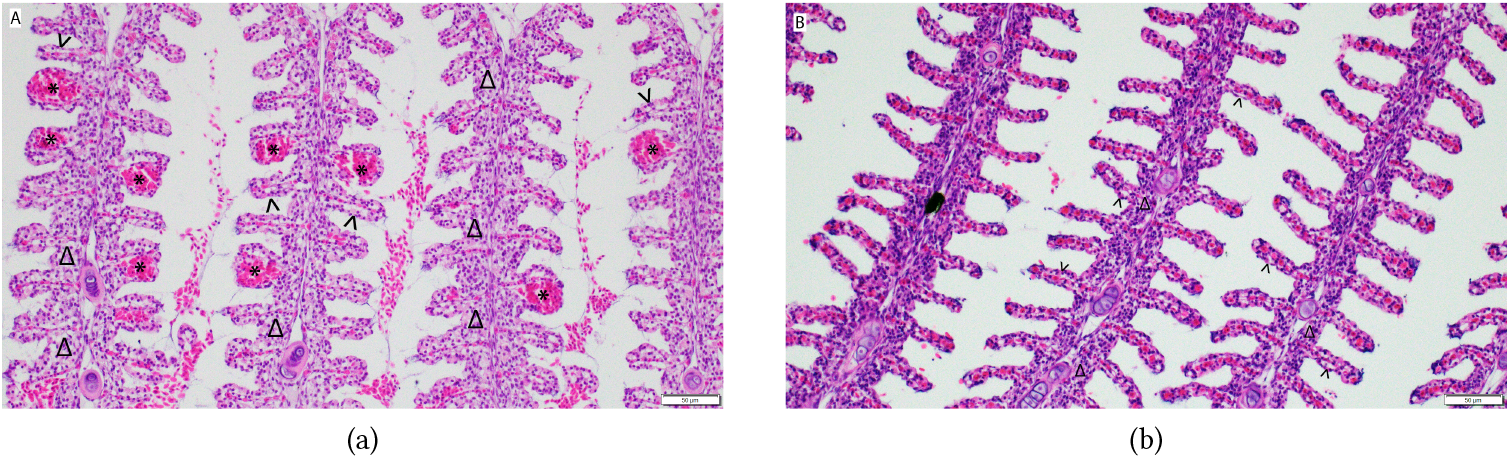
Example of gill lesions observed in three-spot gourami (*Trichopodus trichopterus*) after euthanasia by bath with clove oil or tricaine. (a): Severe gill lesions with moderate to severe aneurisms, moderate epithelial lifting and moderate to severe oedema. (b): Mild gill lesions with no aneurisms, mild epithelial lifting and scattered oedema. Asterisk = aneurisms; arrowhead = epithelial lifting; triangle = oedema.

**Figure 4.**
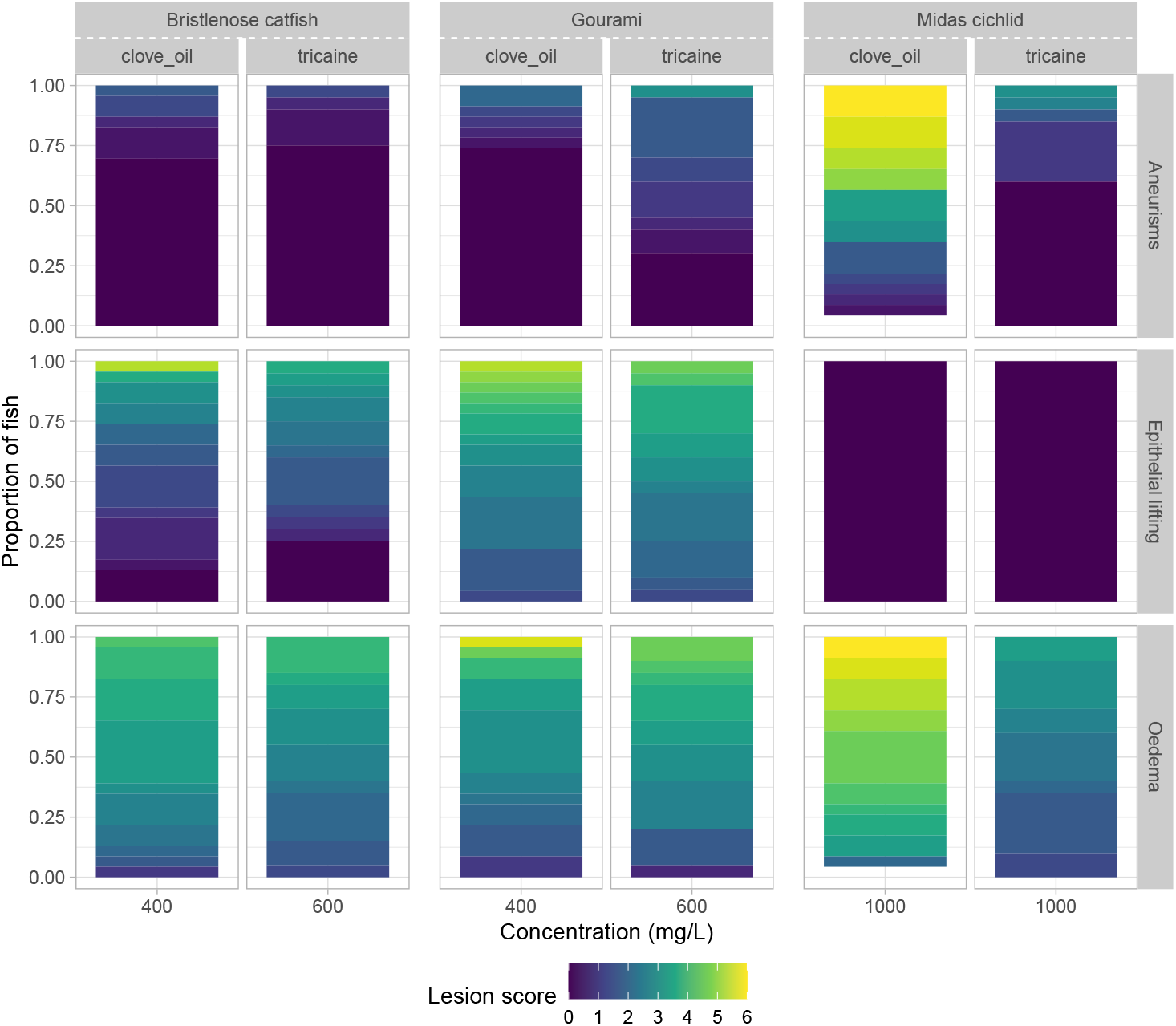
Comparison of lesion scores observed in histological examination for bristlenose catfish (*Ancistrus* sp.), Midas cichlid (*Amphilophus citrinellus*) and three-spot gourami (*Trichopodus trichopterus*) after euthanasia by bath with clove oil and tricaine. Colors represent the severity grade, ranging from dark blue = 0 (no histopathological changes) to yellow = 6 (severe histopathological changes).

Regarding epithelial proliferation, presence of eosinophilic granular cells and presence of parasites, the only significant difference before Bonferroni-Holm correction was for presence of eosinophilic granular cells for three-spot gourami (p = 0.0283). However, this significance was lost following Bonferroni-Holm correction.

## Discussion

### Ability of clove oil to induce loss of indicators in fish

Clove oil did not induce reliable CoOM in goldfish, even at high concentration. However, it provoked loss of all indicators in all other species, with three-spot gourami and bristlenose catfish already losing indicators at lower concentrations (400 mg/L), while Midas cichlids required a higher clove oil concentration (1000 mg/L). In the literature, effective concentrations most often range between 50 and 400 mg/L (Ayala-Soldado et al. 2023; Fernandes et al. 2016; Gneiding et al. 2024; Davis et al. 2015; da Cunha et al. 2015; Ferreira et al. 2022; Holloway et al. 2004; Smith and Harms 2023), with higher concentrations reported for koi carps (*Cyprinus carpio*; > 500 and 1000 mg/L (Gladden et al. 2010; Day et al. 2024)) and Unga cichlids (*Sarotherodon linellii*; 500 mg/L (Bradley et al. 2021)).

Several factors may explain the failure to find a clove oil concentration that reliably induced the loss of all indicators in goldfish and the relatively high concentration required to achieve the endpoints in Midas cichlids. Indeed, goldfish and Midas cichlids are both hypoxiatolerant species, which are assumed to be more difficult to euthanise with a chemical agent and to require a higher dosage than other species (Leary et al. 2020; Balko et al. 2018). The reported concentrations of clove oil found in the literature for euthanasia of other hypoxia-tolerant species, including koi carps and Unga cichlids, are also at the higher end of the reported range (Day et al. 2024; Gladden et al. 2010; Bradley et al. 2021).

However, failure to reliably induce the loss of indicators in goldfish, even at high concentrations of clove oil was unexpected given that previous studies have demonstrated reliable euthanasia in koi carp, a closely related species (Day et al. 2024), and have shown that clove oil can effectively induce anaesthesia in goldfish and in the closely related crucian carp (*Carassius carassius* (Endo et al. 1972)) as well as in Midas cichlids (Gladden et al. 2010). Endo et al. (1972) even described 80 % mortality in goldfish with a concentration of 50 mg/L of eugenol.

A potential explanation for this discrepancy between our study and the literature could be the origin of the fish used. Indeed, the goldfish originated from two private ponds with dense aquatic vegetation and fish overpopulation, conditions commonly associated with low dissolved oxygen concentrations (Wetzel 2001; Boyd and Tucker 1998). Although dissolved oxygen concentrations were not measured, we hypothesize that the goldfish were probably accustomed to severe hypoxic conditions. This could potentially explain the failure of clove oil to reliably induce CoOM and loss of indicators in our goldfish. As reliability is a prerequisite for a suitable euthanasia method, we do not consider eugenol to be a suitable euthanasia agent for goldfish.

### Reliability and interpretation of loss of indicators

Von Krogh et al. (2021) proposed the startle reflex stimulated by tapping at the benchtop surface next to the tank as indicator to assess loss of consciousness during anesthesia. In our experiment, this reflex was harder to test and less informative than assumed, as we incidentally found that we could trigger it in several bristlenose catfish through light finger taps on the table even though the fish did not react to the marble test. We can hypothesize that the intensity of the test using a 76 g marble was too high to trigger the reflex. If startle reflex is needed in a certain context, we therefore recommend to first validate the method that will be used for the species and context of interest, as results could otherwise be biased.

Additionally, it should be noted that the loss of the tested indicators by itself does not necessarily demon-strates a loss of consciousness, defined as a depression of the central nervous system (CNS). However, loss of consciousness is a requisite to attain general anaesthesia (Lamont et al. 2024). This is of particular importance because, although a considerable number of publications claim that clove oil leads to loss of consciousness in fish or has a depressive effect on their central nervous system (Davis et al. 2015; Javahery et al. 2012; Fernandes et al. 2016; Ferreira et al. 2022; Jiang et al. 2023; Soto and Burhanuddin 1995), we could only find four publications that tested it experimentally (Barbas et al. 2021; Wallbom et al. 2026; Machnik et al. 2023; Lehotzky et al. 2023). Only one of these supports this claim. It seems that Hikasa et al. (1986) hypothesized that their results could indicate a depression of the central nervous system with-out testing it, a hypothesis that was then reframed into an accepted claim through citation cascade. Additionally, studies by Ingvast-Larsson et al. (2003) and Barbas et al. (2021) both suggested that clove oil could potentially only induce paralysis and no depression of the central nervous system. However, the experimental set-up of the study by Ingvast-Larsson et al. (2003) was ex-vivo and the experimental set-up of the study by Barbas et al. (2021) aimed solely for anaesthesia, utilising considerably lower clove oil concentrations; therefore, results do not allow for direct interpretation for the context of euthanasia. Machnik et al. (2023) also reported that isoeugenol does not act on central neurons in goldfish in a study that used a lower concentration of isoeugenol, suitable for anaesthesia. However, our results show that goldfish cannot be euthanised reliably with clove oil, potentially indicating that it is not the ideal model species to investigate effects of clove oil compounds on the central nervous system in fish. In addition, Wallbom et al. (2026) reported that neurophysiological measurements of rainbow trout (*Oncorhynchus mykiss*) seem to indicate that rainbow trout lose consciousness when anesthetized with eugenol, but highlighted contradictory results between behavioural and neurophysiological assessments, leaving the question unresolved. On contrary, Lehotzky et al. (2023) showed that eugenol caused a decrease in the frequency of the electrical organ discharge of the brown ghost knifefish (*Apteronotus leptorhynchus*), which is controlled by the pacemaker nucleus situated in the medulla oblongata. This decrease in frequency is, according to the authors, an indication that eugenol can affect the CNS of certain fish species.

Therefore, we suggest prudence when interpreting loss of behavioural indicators in fish, as the relation between loss of indicators and depression of the CNS is currently still unresolved.

### Ability of clove oil to provoke irreversible loss of vital function in fish

None of the three-spot gourami nor of the bristlenose catfish recovered after being immersed in the clove oil solution until the loss of indicators and for an additional 10 minutes thereafter, followed by transfer to water of optimal quality for one hour. These results indicate that clove oil might provoke irreversible loss of vital function in both fish species. However, five out of thirty Midas cichlids did show signs of consciousness within the onehour recovery period, indicating that, in this species, euthanasia attempts with clove oil may not be irreversible and that a secondary killing method is necessary for this species and potentially other hypoxia-tolerant species, as recommended by Leary et al. (2020).

Interestingly, all Midas cichlids that showed recovery signs were on the upper end of the weight distribution. Although the results of a logistic regression found that weight was not a significant predictor for reversibility of euthanasia attempt, we want to point out the fact that only five out of thirty Midas cichlids showed recovery and that weight is not normally distributed in our sample size and includes some high outliers. Rare outcome, imbalance in the outcome size and outliers in predictors can all have effects on the uncertainty, potentially masking significant effects. Therefore, we recommend keeping in mind that weight of Midas cichlids may affect its ability to recover after euthanasia attempt, and that further research is needed to confirm or disprove this hypothesis. In previous studies, association between body size and anaesthetic response was found (Zahl et al. 2009; Skår et al. 2017; Rożyński et al. 2018; Obirikorang et al. 2020), whereas a meta-analysis by Félix et al. (2023) reported no correlation between anesthesia induction and recovery time and fish weight. We therefore recommend to use a secondary killing method when euthanising large fish.

However, since the available data is still limited, and to apply precautionary principles to avoid unnecessary suffering of the fish, we generally recommend using a secondary killing method to ensure death after induction with clove oil, as recommended by Leary et al. (2020), regardless of the species and size of the fish.

### Suitability of clove oil for humane euthanasia of fish

Leary et al. (2020) states that the selected euthanasia method “should result in rapid loss of consciousness followed by cardiac or respiratory arrest and, ultimately, a loss of brain function. In addition, animal handling and the euthanasia technique should minimize distress experienced by the animal prior to loss of consciousness.” Therefore, treatments that minimize behavioural adverse reactions and histological lesions would be more appropriate, as distress would thus be minimized. The histology results show that Midas cichlids have significantly more lesions when euthanised with clove oil than when euthanised with tricaine. It is noteworthy to remark that not only was the difference significant, but also that the Hodges–Lehmann estimate of the median shift was substantial. Based on these results, we suggest that clove oil is not adapted for the euthanasia of Midas cichlids. Behavioural observations also suggest a potential adverse reaction to clove oil, as all Midas cichlids euthanised with clove oil expressed forceful buccal pumping, which was not observed for Midas cichlids exposed to the negative control. However, Midas cichlids euthanised with tricaine also expressed forceful buccal pumping, and Midas cichlids showed longer and more intense episodes of fast swimming when euthanised with tricaine than with clove oil. These results suggest than none of the substances are optimal to euthanise Midas cichlids, possibly because the dosage needed to achieve euthanasia are particularly high, leading to adverse reactions.

In contrary, three-spot gourami had significantly less lesions with clove oil than with tricaine, although the Hodges–Lehmann estimate of the median shift was far smaller. However, only one concentration of tricaine was tested and therefore no step was taken beforehand to optimize the concentration, as was performed for clove oil. However, six out of twenty three-spot gourami euthanised with clove oil recovered movement after having lost both movement and righting reflex. Although these movements were then lost again, we cannot exclude that these episodes are stressful to the fish. Therefore, considering the small Hodges–Lehmann estimate of the median shift for the score of the histological lesions, the lack of optimization for tricaine concentration, and the behavioural observations, we do not consider these results to be clear enough and thus refrain from making recommendations on the best substance for the euthanasia of threespot gourami. Similarly, we cannot make substance recommendations for the euthanasia of bristlenose gourami, as differences were found neither in behavioural observations nor in histology.

Additionally, we already mentioned above the lack of experimental proof that clove oil exposure induces a CNS depression in fish and therefore leads to loss of consciousness. This directly questions the overall suitability of clove oil as an agent to perform humane euthanasia, at least until this question is resolved.

## Conclusion and animal welfare implications

In conclusion, we showed that clove oil is not suitable for humane euthanasia of goldfish and Midas cichlids, and postulate that these findings are likely to be valid for other hypoxia-tolerant species. We also highlighted that questions regarding the suitability of clove oil to euthanise other fish species are not resolved, since the ability of clove oil to induce loss of consciousness and not only paralysis in fish has not been demonstrated yet. Therefore, we currently recommend avoiding euthanising fish with clove oil when alternatives such as tricaine are available. In all cases, we also recommend using a secondary killing method as recommended by Leary et al. (2020), as we showed that some fish could recover after euthanasia attempt with clove oil.

## Acknowledgements

We thank the Papiliorama Foundation in Kerzers for its collaboration and support with the study involving bristlenose catfish and Midas cichlids. We further thank the histology technicians at the Institute of Veterinary Pathology, University of Bern for their help in laboratory work. We also thank Manuel Blonç from the Institute for Fish and Wildlife Health, University of Bern for his valuable proofreading of the manuscript.

## Appendix

**Appendix 1.**
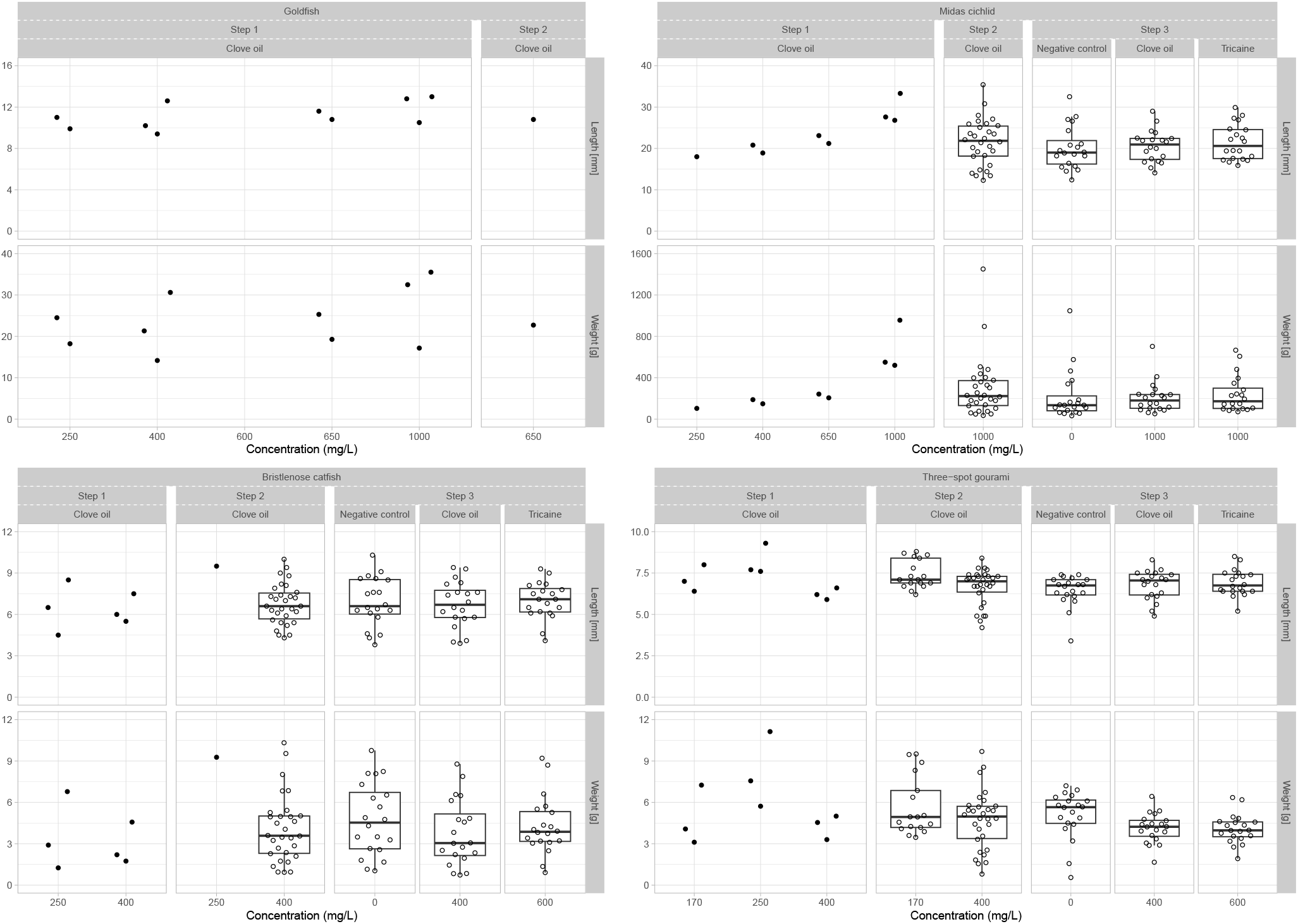
Distribution of fish size and weight for each ental step combination of species, treatment, concentration and experimental step

**Appendix 2.**
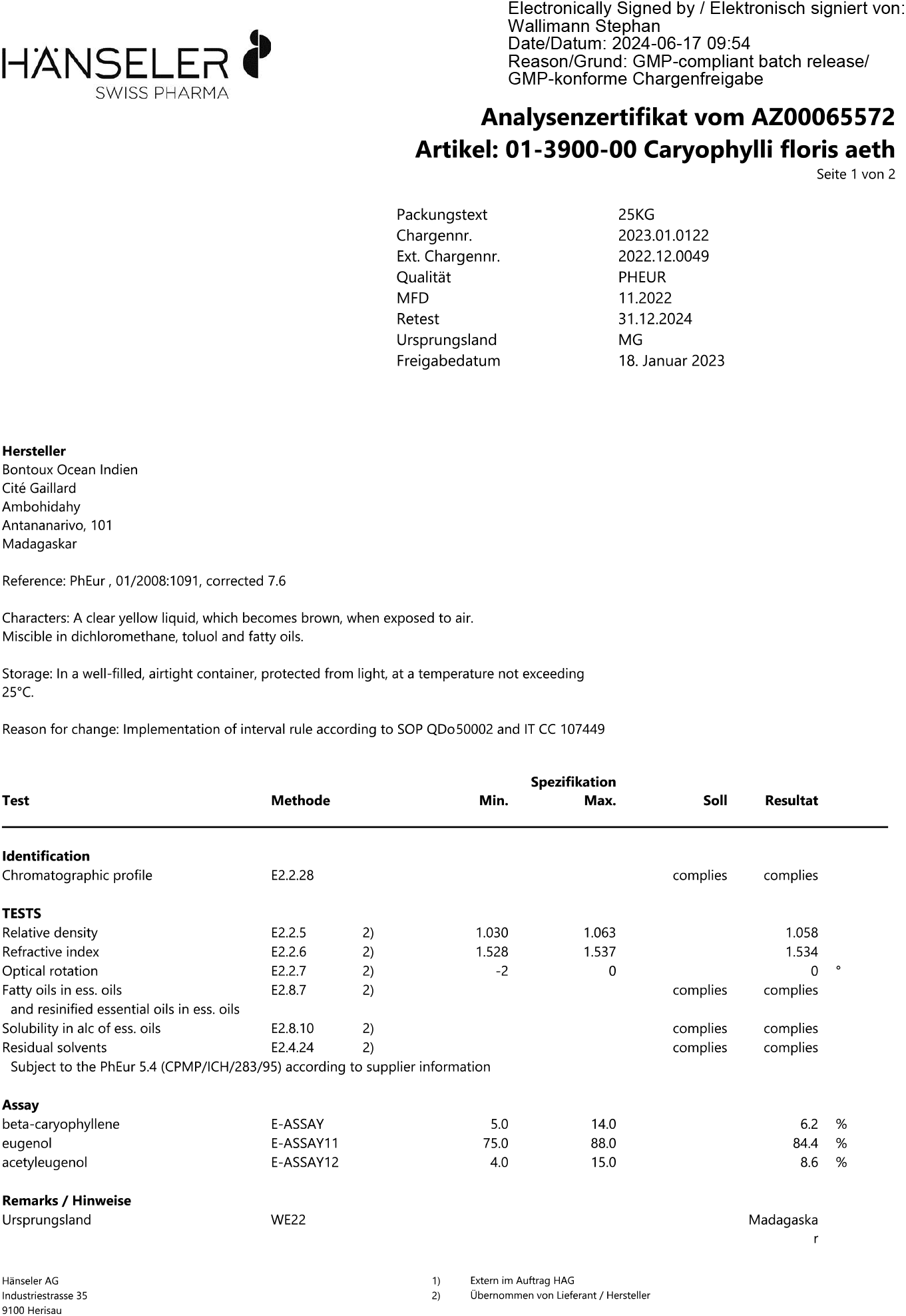
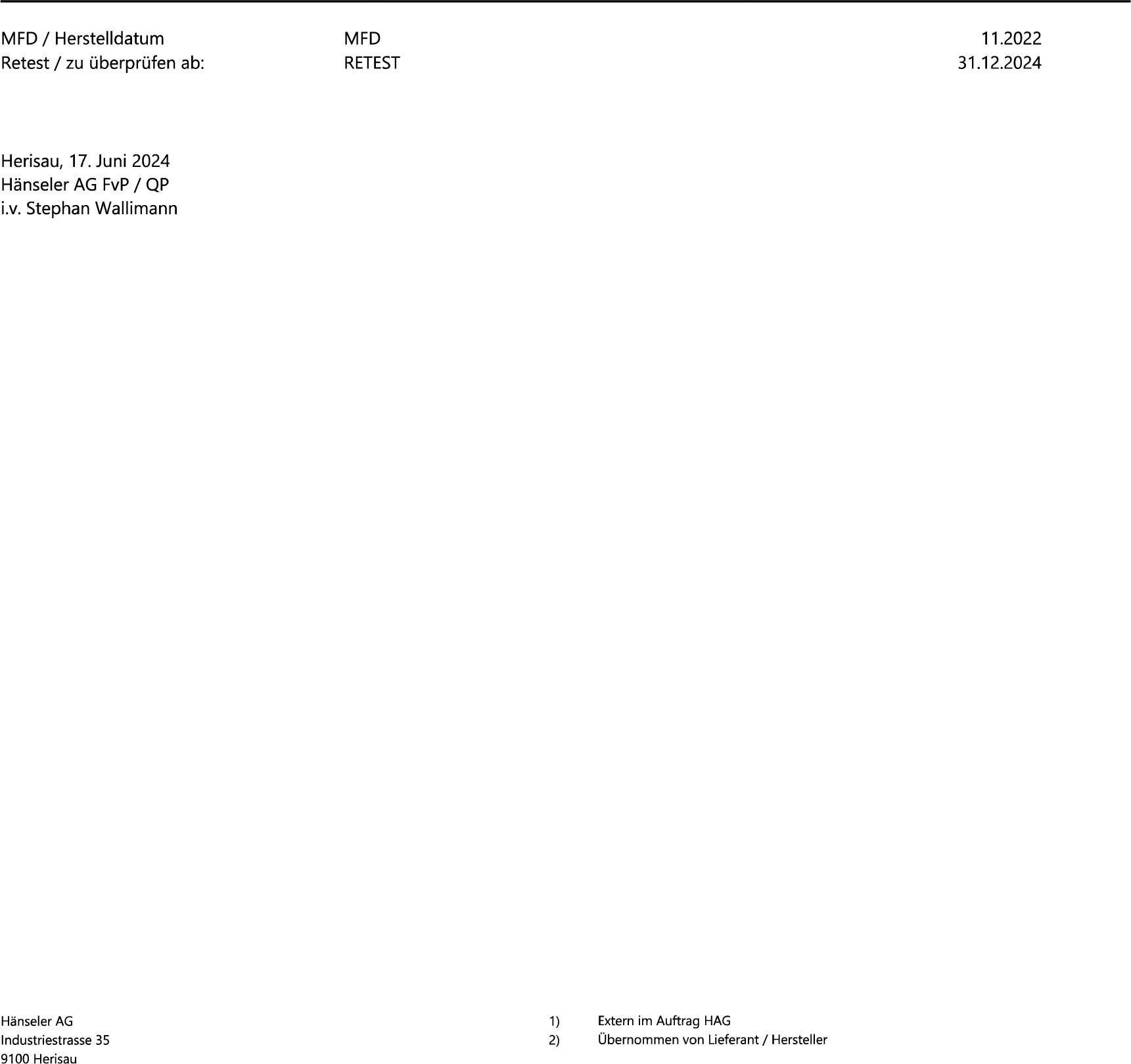
Certificate of analysis of the clove oil used in the study

**Appendix 3.** Time to loss of indicators for goldfish (*Carassius auratus*), bristlenose catfish (*Ancistrus* sp.), Midas cichlid (*Amphilophus citrinellus*) and three-spot gourami (*Trichopodus trichopterus*) treated with different concentration of eugenol diluted in the water. Additionally, fish size and weight as well as water temperature are provided. LoRR: loss of righting reflex / CoOM: cessation of opercular movement / LoSR: loss of startle reflex

| Concentration [mg/L] | Weight [g] | Length [cm] | Temperature [°C] | LoRR | Time to LoRR [s] | CoOM | Time to CoOM [s] | LoSR | Time to LoSR [s] |
|---|---|---|---|---|---|---|---|---|---|
| Goldfish |  |  |  |  |  |  |  |  |  |
| 250 | 18.22 | 9.9 | 17.4 | yes | 40 | no | NA | absent | NA |
|  | 24.51 | 11.0 | 17.5 | yes | 60 | yes | 1470 | absent | NA |
| 400 | 14.17 | 9.4 | 18.2 | yes | 31 | yes | 449 | absent | NA |
|  | 30.62 | 12.6 | 18.6 | yes | 32 | no | NA | absent | NA |
|  | 21.32 | 10.2 | 18.4 | yes | 52 | yes | 469 | absent | NA |
| 650 | 19.28 | 10.8 | 18.5 | yes | 44 | yes | 456 | absent | NA |
|  | 25.31 | 11.6 | 18.9 | yes | 38 | yes | 933 | absent | NA |
|  | 22.72 | 10.8 | 18.2 | yes | 31 | no | NA | yes | 75 |
| 1000 | 32.49 | 12.8 | 14.2 | yes | 36 | yes | 271 | yes | 150 |
|  | 17.17 | 10.5 | 15.0 | yes | 61 | yes | 261 | yes | 15 |
|  | 35.52 | 13.0 | 15.7 | yes | 62 | no | NA | yes | 75 |
| Midas cichlid |  |  |  |  |  |  |  |  |  |
| 250 | 103.70 | 18.0 | 20.5 | yes | 37 | no | NA | absent | NA |
| 400 | 148.37 | 18.9 | 20.5 | yes | 33 | yes | 811 | absent | NA |
|  | 188.10 | 20.8 | 19.9 | yes | 24 | no | NA | absent | NA |
| 650 | 241.94 | 23.1 | 20.3 | yes | 23 | yes | 15 | absent | NA |
|  | 205.93 | 21.2 | 20.2 | yes | 21 | no | NA | absent | NA |
| 1000 | 520.01 | 26.8 | 21.2 | yes | 29 | yes | 29 | absent | NA |
|  | 550.35 | 27.6 | 20.9 | yes | 33 | yes | 33 | absent | NA |
|  | 956.21 | 33.3 | 20.8 | yes | 43 | yes | 61 | absent | NA |
|  | 179.27 | 18.3 | 22.5 | yes | 16 | yes | 16 | yes | 15 |
|  | 230.40 | 22.1 | 22.4 | yes | 18 | yes | 18 | absent | NA |
| 1000 | 48.56 | 13.4 | 22.3 | yes | 19 | yes | 19 | absent | NA |
|  | 328.60 | 24.0 | 22.3 | yes | 22 | yes | 22 | absent | NA |
|  | 49.91 | 13.4 | 22.3 | yes | 16 | yes | 21 | yes | 14 |
|  | 180.10 | 20.2 | 22.3 | yes | 16 | yes | 31 | absent | NA |
|  | 103.98 | 14.0 | 22.2 | yes | 28 | yes | 22 | yes | 18 |
|  | 35.30 | 12.3 | 22.2 | yes | 18 | yes | 19 | absent | NA |
|  | 1451.21 | 35.4 | 21.7 | yes | 26 | yes | 18 | absent | NA |
|  | 376.10 | 25.9 | 21.7 | yes | 25 | yes | 18 | absent | NA |
|  | 60.10 | 14.4 | 22.6 | yes | 17 | yes | 19 | yes | 15 |
|  | 254.20 | 22.4 | 22.6 | yes | 22 | yes | 13 | absent | NA |
|  | 129.00 | 18.1 | 22.6 | yes | 27 | yes | 27 | yes | 25 |
|  | 202.00 | 21.4 | 22.6 | yes | 28 | yes | 21 | absent | NA |
|  | 229.20 | 21.6 | 22.4 | yes | 26 | yes | 20 | absent | NA |
|  | 398.70 | 26.0 | 22.1 | yes | 22 | yes | 14 | absent | NA |
|  | 438.12 | 26.6 | 22.1 | yes | 30 | yes | 12 | absent | NA |
|  | 75.14 | 15.9 | 22.0 | yes | 19 | yes | 22 | absent | NA |
|  | 76.34 | 14.8 | 22.0 | yes | 18 | yes | 18 | absent | NA |
|  | 506.10 | 28.0 | 22.0 | yes | 30 | yes | 20 | yes | 10 |
|  | 402.40 | 25.1 | 22.4 | yes | 30 | yes | 23 | yes | 10 |
|  | 302.90 | 23.6 | 22.3 | yes | 22 | yes | 11 | absent | NA |
|  | 317.70 | 23.0 | 22.3 | yes | 31 | yes | 21 | absent | NA |
|  | 194.30 | 20.5 | 22.4 | yes | 16 | yes | 14 | absent | NA |
|  | 216.30 | 23.5 | 22.2 | yes | 42 | yes | 18 | absent | NA |
|  | 896.00 | 30.8 | 22.0 | yes | 120 | yes | 15 | absent | NA |
|  | 360.60 | 25.5 | 22.0 | yes | 30 | yes | 16 | absent | NA |
|  | 480.10 | 27.1 | 21.9 | yes | 24 | yes | 13 | absent | NA |
| 1000 | 159.40 | 19.4 | 21.9 | yes | 16 | yes | 14 | absent | NA |
|  | 138.10 | 19.2 | 21.9 | yes | 29 | yes | 29 | yes | 17 |
|  | 236.70 | 22.1 | 22.8 | yes | 21 | yes | 26 | absent | NA |
|  | 326.51 | 24.2 | 22.8 | yes | 19 | yes | 25 | absent | NA |
|  | 134.70 | 19.3 | 22.7 | yes | 22 | yes | 14 | absent | NA |
|  | 153.46 | 20.0 | 22.7 | yes | 21 | yes | 18 | absent | NA |
|  | 288.80 | 23.8 | 22.7 | yes | 14 | yes | 64 | absent | NA |
|  | 410.60 | 26.6 | 22.7 | yes | 31 | yes | 30 | absent | NA |
|  | 228.30 | 21.9 | 22.7 | yes | 24 | yes | 24 | absent | NA |
|  | 207.00 | 22.5 | 22.6 | yes | 17 | yes | 20 | absent | NA |
|  | 207.80 | 21.6 | 22.6 | yes | 18 | yes | 18 | yes | 15 |
|  | 100.20 | 17.5 | 22.6 | yes | 27 | yes | 30 | absent | NA |
|  | 240.02 | 22.4 | 22.8 | yes | 24 | yes | 26 | absent | NA |
|  | 89.88 | 16.7 | 22.8 | yes | 27 | yes | 30 | yes | 17 |
|  | 702.90 | 29.0 | 22.8 | yes | 23 | yes | 23 | absent | NA |
|  | 62.67 | 15.3 | 22.6 | yes | 17 | yes | 11 | absent | NA |
|  | 236.65 | 22.0 | 22.6 | yes | 25 | yes | 27 | absent | NA |
|  | 86.67 | 16.9 | 22.7 | yes | 25 | yes | 14 | absent | NA |
|  | 151.66 | 20.3 | 22.7 | yes | 24 | yes | 12 | absent | NA |
|  | 106.22 | 18.1 | 22.7 | yes | 19 | yes | 18 | absent | NA |
|  | 116.23 | 16.5 | 22.6 | yes | 23 | yes | 16 | absent | NA |
|  | 51.77 | 14.1 | 22.5 | yes | 15 | yes | 15 | yes | 15 |
| Bristlenose catfish |  |  |  |  |  |  |  |  |  |
| 250 | 6.78 | 8.5 | 23.2 | yes | 36 | yes | 500 | absent | NA |
|  | 2.90 | 6.5 | 22.2 | yes | 80 | yes | 504 | absent | 110 |
|  | 1.25 | 4.5 | 21.7 | yes | 52 | yes | 270 | absent | NA |
| 250 | 9.27 | 9.5 | 22.9 | yes | 79 | no | NA | absent | NA |
| 400 | 2.20 | 6.0 | 21.1 | yes | 52 | yes | 260 | absent | NA |
|  | 1.74 | 5.5 | 20.5 | yes | 46 | yes | 232 | absent | NA |
|  | 4.57 | 7.5 | 20.3 | yes | 49 | yes | 270 | absent | NA |
|  | 8.02 | 9.0 | 23.1 | yes | 100 | yes | 480 | absent | NA |
|  | 0.95 | 4.5 | 23.2 | yes | 29 | yes | 180 | absent | NA |
|  | 1.35 | 4.8 | 22.4 | yes | 45 | yes | 240 | absent | NA |
|  | 6.85 | 8.8 | 21.8 | yes | 76 | yes | 840 | yes | 120 |
|  | 10.32 | 10.0 | 23.6 | yes | 66 | yes | 198 | yes | 198 |
|  | 3.42 | 6.2 | 23.9 | yes | 52 | yes | 597 | absent | NA |
|  | 0.95 | 4.5 | 24.1 | yes | 53 | yes | 360 | yes | 88 |
|  | 4.95 | 8.2 | 23.1 | yes | 50 | yes | 563 | yes | 143 |
|  | 3.60 | 6.6 | 23.0 | yes | 99 | yes | 665 | absent | NA |
|  | 2.68 | 6.4 | 22.3 | yes | 54 | yes | 430 | yes | 172 |
|  | 9.54 | 9.4 | 23.5 | yes | 63 | yes | 603 | yes | 223 |
|  | 6.83 | 8.1 | 23.2 | yes | 57 | yes | 614 | yes | 103 |
|  | 4.98 | 7.4 | 22.6 | yes | 53 | yes | 350 | yes | 134 |
|  | 2.28 | 5.6 | 21.9 | yes | 48 | yes | 421 | yes | 140 |
|  | 5.26 | 7.3 | 23.3 | yes | 103 | yes | 395 | yes | 162 |
|  | 4.96 | 7.3 | 24.2 | yes | 69 | yes | 302 | yes | 92 |
|  | 4.05 | 7.0 | 23.8 | yes | 50 | yes | 252 | yes | 65 |
|  | 2.85 | 6.4 | 23.4 | yes | 50 | yes | 405 | yes | 70 |
|  | 3.47 | 6.6 | 23.1 | yes | 62 | yes | 324 | yes | 140 |
|  | 0.93 | 4.3 | 22.5 | yes | 56 | yes | 219 | yes | 99 |
|  | 1.74 | 5.4 | 22.3 | yes | 67 | yes | 271 | yes | 98 |
|  | 5.42 | 7.9 | 24.3 | yes | 87 | yes | 468 | yes | 118 |
| 400 | 4.49 | 7.6 | 23.9 | yes | 97 | yes | 336 | yes | 202 |
|  | 4.62 | 7.1 | 22.9 | yes | 52 | yes | 261 | yes | 94 |
|  | 2.36 | 5.4 | 22.7 | yes | 53 | yes | 227 | absent | NA |
|  | 3.24 | 6.1 | 24.3 | yes | 41 | yes | 255 | yes | 58 |
|  | 1.67 | 5.2 | 24.1 | yes | 36 | yes | 274 | absent | NA |
|  | 5.02 | 7.2 | 23.6 | yes | 61 | yes | 490 | yes | 117 |
|  | 2.09 | 5.9 | 23.2 | yes | 52 | yes | 440 | yes | 79 |
|  | 3.56 | 6.3 | 22.9 | yes | 63 | yes | 378 | yes | 75 |
|  | 0.73 | 3.9 | 24.3 | yes | 31 | yes | 250 | absent | NA |
|  | 7.88 | 9.3 | 23.9 | yes | 32 | yes | 732 | absent | NA |
|  | 6.49 | 8.2 | 23.5 | yes | 35 | yes | 631 | absent | NA |
|  | 3.06 | 6.5 | 23.2 | yes | 69 | yes | 316 | absent | NA |
|  | 0.84 | 4.0 | 22.9 | yes | 44 | yes | 203 | absent | NA |
|  | 6.13 | 8.7 | 24.5 | yes | 92 | yes | 381 | absent | NA |
|  | 3.03 | 6.9 | 24.2 | yes | 56 | yes | 445 | absent | NA |
|  | 3.82 | 7.6 | 23.8 | yes | 52 | yes | 420 | absent | NA |
|  | 2.21 | 5.7 | 23.5 | yes | 120 | yes | 374 | absent | NA |
|  | 2.34 | 6.2 | 23.3 | yes | 55 | yes | NA | absent | NA |
|  | 4.57 | 7.5 | 24.4 | yes | 57 | yes | NA | absent | 575 |
|  | 1.45 | 5.1 | 23.8 | yes | 43 | yes | NA | absent | NA |
|  | 0.83 | 4.1 | 23.1 | yes | 53 | yes | 512 | absent | 626 |
|  | 1.95 | 5.8 | 22.8 | yes | 28 | yes | 256 | absent | 536 |
|  | 4.75 | 7.6 | 22.5 | yes | 45 | yes | 425 | absent | NA |
|  | 6.57 | 8.3 | 23.9 | yes | 45 | yes | 326 | absent | 559 |
|  | 2.52 | 6.3 | 23.4 | yes | 38 | yes | 260 | absent | 348 |
|  | 4.84 | 7.4 | 23.1 | yes | 97 | yes | 348 | absent | 391 |
| 400 | 2.75 | 5.8 | 22.8 | yes | 56 | yes | 240 | absent | 500 |
|  | 8.78 | 9.4 | 22.6 | yes | 45 | yes | 492 | absent | 660 |
| Three-spot gourami |  |  |  |  |  |  |  |  |  |
| 170 | 4.07 | 7.0 | 24.6 | yes | 116 | yes | 405 | absent | NA |
|  | 3.11 | 6.4 | 23.7 | yes | 86 | yes | 357 | absent | NA |
|  | 7.25 | 8.0 | 22.9 | yes | 46 | yes | 361 | absent | NA |
|  | 4.56 | 7.1 | 22.4 | yes | 64 | yes | 573 | absent | NA |
|  | 3.46 | 6.2 | 24.6 | yes | 31 | yes | 429 | absent | 53 |
|  | 6.37 | 7.8 | 23.7 | yes | 52 | yes | 352 | absent | NA |
|  | 4.25 | 7.3 | 23.5 | yes | 42 | yes | 266 | absent | 61 |
|  | 8.90 | 8.5 | 24.1 | yes | 44 | yes | 479 | absent | NA |
|  | 8.32 | 8.6 | 23.6 | yes | 45 | yes | 430 | absent | NA |
|  | 4.95 | 7.3 | 23.3 | yes | 48 | yes | 780 | absent | 68 |
|  | 4.09 | 7.1 | 24.3 | yes | 72 | yes | 681 | absent | NA |
|  | 4.18 | 6.9 | 23.8 | yes | 42 | yes | 600 | absent | NA |
|  | 5.05 | 6.7 | 24.1 | yes | 66 | yes | 607 | absent | NA |
|  | 9.47 | 8.8 | 23.6 | yes | 61 | yes | 570 | absent | NA |
|  | 3.88 | 6.4 | 24.1 | yes | 319 | yes | 489 | absent | NA |
|  | 9.51 | 8.7 | 23.1 | yes | 45 | yes | 635 | absent | NA |
|  | 6.86 | 8.4 | 23.5 | yes | 32 | yes | 565 | absent | NA |
|  | 4.43 | 6.9 | 23.1 | yes | 55 | yes | 750 | absent | NA |
|  | 4.94 | 6.9 | 22.4 | yes | 225 | yes | 720 | absent | 270 |
|  | 3.60 | 6.7 | 23.3 | yes | 123 | yes | 455 | absent | NA |
| 250 | 7.56 | 7.7 | 24.9 | yes | 56 | yes | 497 | absent | NA |
|  | 11.13 | 9.3 | 24.1 | yes | 33 | yes | 568 | absent | NA |
|  | 5.72 | 7.6 | 23.8 | yes | 59 | yes | 500 | absent | NA |
| 400 | 4.54 | 6.2 | 24.9 | yes | 33 | yes | 33 | absent | NA |
|  | 3.30 | 5.9 | 24.6 | yes | 43 | yes | 423 | absent | NA |
|  | 5.00 | 6.6 | 24.2 | yes | 21 | yes | 509 | absent | NA |
|  | 4.80 | 6.7 | 24.8 | yes | 40 | yes | 353 | absent | NA |
|  | 6.36 | 7.3 | 24.3 | yes | 29 | yes | 555 | absent | NA |
|  | 6.14 | 7.3 | 24.1 | yes | 25 | yes | 457 | absent | NA |
|  | 4.83 | 6.8 | 23.8 | yes | 25 | yes | 489 | absent | NA |
|  | 4.42 | 6.7 | 23.5 | yes | 37 | yes | 189 | absent | NA |
|  | 6.72 | 7.7 | 23.2 | yes | 36 | yes | 393 | absent | NA |
|  | 5.10 | 7.1 | 23.1 | yes | 28 | yes | 569 | absent | NA |
|  | 5.40 | 6.9 | 22.8 | yes | 19 | yes | 342 | absent | NA |
|  | 5.43 | 7.2 | 22.6 | yes | 40 | yes | 492 | absent | NA |
|  | 3.55 | 6.3 | 23.9 | yes | 78 | yes | 382 | absent | NA |
|  | 5.78 | 7.3 | 23.6 | yes | 38 | yes | 537 | absent | NA |
|  | 5.37 | 7.1 | 23.0 | yes | 68 | yes | 374 | absent | NA |
|  | 1.82 | 4.9 | 22.8 | yes | 31 | yes | 389 | absent | NA |
|  | 4.06 | 6.8 | 22.5 | yes | 33 | yes | 452 | absent | NA |
|  | 4.84 | 7.4 | 22.1 | yes | 66 | yes | 247 | absent | NA |
|  | 2.54 | 5.7 | 21.9 | yes | 20 | yes | 456 | absent | NA |
|  | 5.18 | 7.4 | 24.0 | yes | 39 | yes | 416 | absent | NA |
|  | 5.67 | 7.2 | 23.9 | yes | 554 | yes | 416 | absent | NA |
|  | 0.80 | 4.2 | 23.3 | yes | 40 | yes | 257 | absent | 55 |
|  | 9.69 | 8.4 | 24.8 | yes | 34 | yes | 250 | absent | NA |
|  | 4.40 | 6.7 | 24.5 | yes | 38 | yes | 285 | absent | NA |
|  | 5.73 | 7.2 | 23.8 | yes | 33 | yes | 148 | absent | NA |
|  | 8.55 | 7.8 | 23.7 | yes | 61 | yes | 435 | absent | NA |
| 400 | 8.17 | 7.8 | 23.4 | yes | 52 | yes | 259 | absent | NA |
|  | 3.32 | 6.5 | 23.1 | yes | 44 | yes | 62 | absent | NA |
|  | 5.48 | 7.4 | 22.8 | yes | 36 | yes | 567 | absent | NA |
|  | 2.38 | 5.4 | 22.5 | yes | 32 | yes | 365 | absent | NA |
|  | 1.63 | 4.9 | 22.3 | yes | 27 | yes | 338 | absent | NA |
|  | 1.55 | 4.6 | 22.0 | yes | 71 | yes | 382 | absent | NA |
|  | 2.19 | 4.9 | 22.0 | yes | 36 | yes | 435 | absent | NA |
|  | 5.18 | 7.6 | 24.7 | yes | 40 | yes | 155 | absent | 115 |
|  | 2.90 | 5.2 | 24.3 | yes | 29 | yes | 543 | absent | NA |
|  | 1.66 | 4.9 | 23.8 | yes | 81 | yes | 512 | absent | NA |
|  | 3.05 | 6.1 | 23.4 | yes | 35 | yes | 35 | absent | NA |
|  | 3.60 | 6.2 | 23.1 | yes | 39 | yes | 480 | absent | 350 |
|  | 3.35 | 6.0 | 24.9 | yes | 35 | yes | 914 | absent | NA |
|  | 2.88 | 5.6 | 24.4 | yes | 79 | yes | 394 | absent | NA |
|  | 5.38 | 7.4 | 24.0 | yes | 24 | yes | 337 | absent | NA |
|  | 4.77 | 7.5 | 23.7 | yes | 45 | yes | 317 | absent | NA |
|  | 4.67 | 7.7 | 23.4 | yes | 52 | yes | 284 | absent | NA |
|  | 4.42 | 7.1 | 25.0 | yes | 42 | yes | 360 | absent | NA |
|  | 4.00 | 6.8 | 24.4 | yes | 24 | yes | 336 | absent | NA |
|  | 5.30 | 7.2 | 23.9 | yes | 31 | yes | 259 | absent | NA |
|  | 4.39 | 7.0 | 23.6 | yes | 31 | yes | 328 | absent | 38 |
|  | 4.24 | 7.3 | 23.2 | yes | 41 | yes | 324 | absent | NA |
|  | 4.45 | 7.5 | 24.7 | yes | 32 | yes | 295 | absent | NA |
|  | 4.22 | 7.1 | 24.3 | yes | 47 | yes | 322 | absent | NA |
|  | 6.43 | 8.3 | 23.9 | yes | 44 | yes | 309 | absent | NA |
|  | 3.86 | 6.3 | 23.6 | yes | 26 | yes | 275 | absent | NA |
| 400 | 3.92 | 6.8 | 23.2 | yes | 37 | yes | 288 | absent | NA |

